# Evidence of phase resetting, not just pulses of sound, during eye movement-related eardrum oscillations (EMREOs)

**DOI:** 10.64898/2026.03.25.714060

**Authors:** Cynthia D King, Christopher A. Shera, Jennifer M Groh

## Abstract

Eye movement-related eardrum oscillations (EMREOs) appear to consist of a pulse of oscillation occurring in conjunction with saccades. However, this apparent pulse could occur either because there is an increase in energy at that frequency at the time of saccades (a true pulse), or because there is saccade-related phase resetting of ongoing energy at that frequency band, thus appearing like a pulse when averaged in the time domain across many trials. Here we conducted a spectral analysis at the individual trial level in humans performing a visually guided saccade task to determine whether the power at the EMREO frequency (30-45 Hz) is higher during saccades than during steady fixation. Although all subjects had easily identifiable EMREOs, we found that only 40% of participants showed a statistically significant increase in sound power in the EMREO frequency band associated with saccade onset, i.e. sound pulses at the individual trial level (in some cases, these power increases were sustained during eccentric fixation). In contrast, 97% of subjects showed statistically significant phase resetting at saccade onset. While both factors may contribute to the apparently pulse-like EMREO signal, phase resetting appears to be more reliably associated with EMREOs. The prevalence of phase resetting has implications for the underlying mechanism(s) producing EMREOs as well as functional consequences for how the ear might respond to incoming sound in an eye-position dependent fashion.

## BACKGROUND

Information about eye movements is necessary for integrating auditory and visual spatial information. Sound location is computed based on cues anchored to the head, whereas visual location is assessed by the pattern of light on the retina. In species with mobile eyes with respect to the head (humans, monkeys, and many other species), the angle of gaze governs the relationship between retinal and head-centered locations. Where and how eye movement signals are incorporated into auditory processing has therefore been of considerable interest for decades (e.g. (Caruso, Pages, Sommer, & Groh, 2021; J.M. Groh, 2014; J. M. Groh & Sparks, 1992; J. M. Groh, Trause, Underhill, Clark, & Inati, 2001; Jay & Sparks, 1984, 1987; Stricanne, Andersen, & Mazzoni, 1996; Zwiers, Versnel, & Van Opstal, 2004)).

Recently, an eye movement signal has been discovered in the auditory periphery (Gruters et al., 2018). Small microphones placed in the ear canals can detect these eye movement-related eardrum oscillations (EMREOs) (Abbasi et al., 2025; Brohl & Kayser, 2023; Gruters et al., 2018; C. D. King et al., 2023; Cynthia D King, Zhu, & Groh, 2026; Leon, Ramos, & Maldonado, 2026; Lovich et al., 2023; Sotero Silva, Brohl, & Kayser, 2026; Sotero Silva, Kayser, & Brohl, 2025). EMREOs can be considered to be a unique form of otoacoustic emissions (OAEs), occurring in association with saccadic eye movements rather than being triggered by external sounds like most other kinds of OAEs. EMREOs occur at a low frequency, ∼30-40Hz, and their amplitude and phase change with saccade magnitude and direction, respectively.

Why EMREOs are oscillatory is a mystery. If EMREOs reflect an underlying mechanism that serves to adjust interaural timing and level differences or spectral cues in an eye position dependent fashion to help prepare for subsequent integration of sounds with visual information (Cho, Ravicz, & Puria, 2023), it is unclear how an oscillatory signal would be beneficial to that process. But do EMREOs in fact consist of pulses of oscillation? They certainly do in averaged recordings in which the individual trials are aligned to saccade onset or offset (Gruters et al., 2018; Lovich et al., 2023). But there are two ways these averaged oscillations could appear. One possibility is that a true pulse of oscillatory power at that frequency occurs in conjunction with each saccade (Figure 1a). Under this scenario, the average of many trials is similar to each individual trial, just less noisy. The second possibility is that there is ongoing power at that frequency all the time, but at the time of the saccade, something happens to reset the phase, bringing the ongoing signal into temporal alignment across trials, and producing the appearance of a pulse of sound in the across-trials average (Figure 1b). These possibilities are not mutually exclusive – either or both could account for EMREOs.

**Figure 1:**
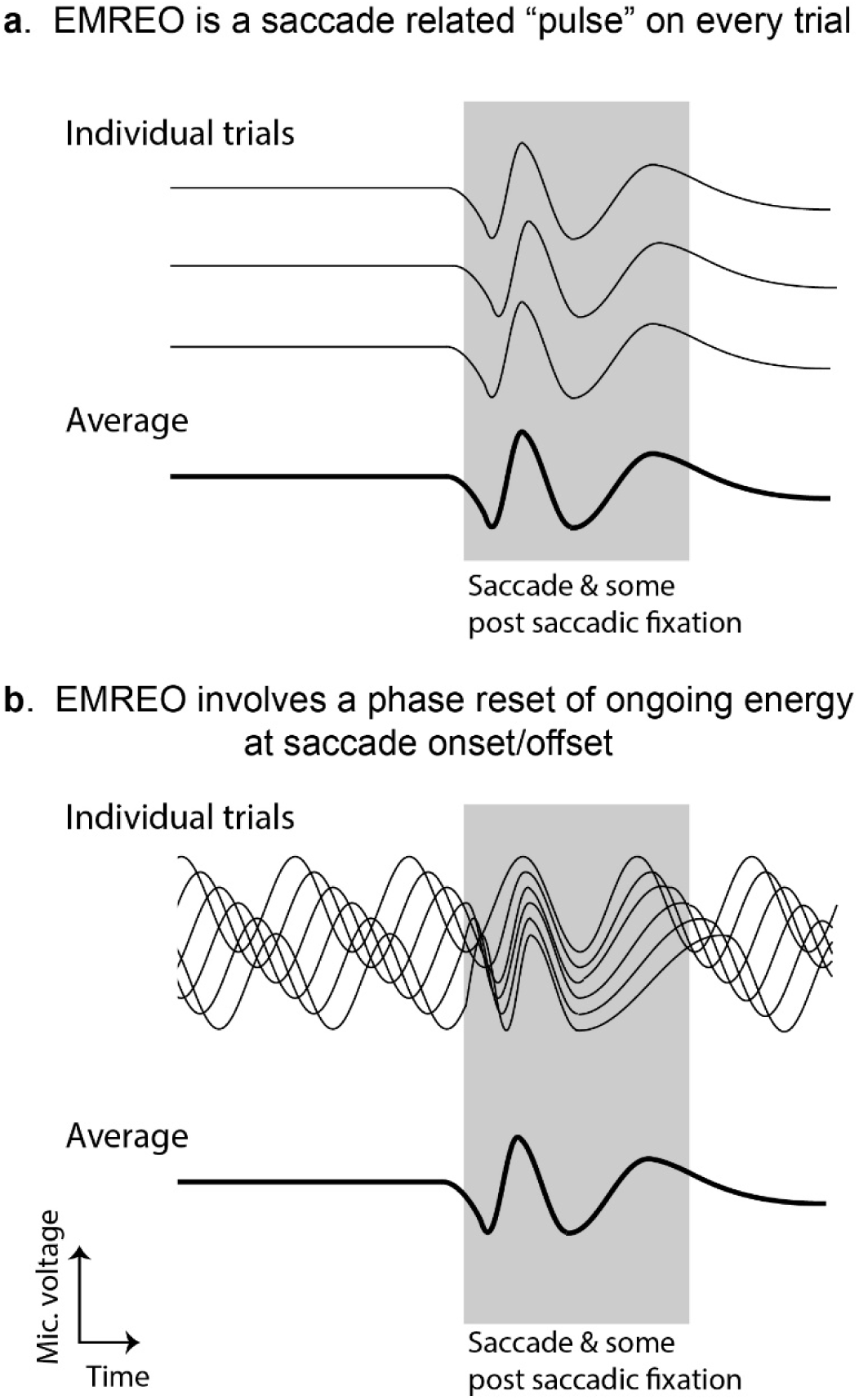
Two competing hypotheses regarding EMREOs. **a.** One possibility is that the EMREO consists of a low frequency “pulse” at about ∼35-40 Hz occurring at the time of the saccade, and that it is evident on every trial in comparison to a flat baseline. Under this scenario, the average trace is a good representation of what is happening on individual trials, and reduces noise with averaging. b. The second possibility is that there is ongoing energy at ∼35-40 Hz even during steady fixation, and that what happens at the time of a saccade is not an increase in that energy but a phase reset, caused by some unknown mechanism. Under this scenario, the EMREO energy would be relatively consistent across time in individual trials, and the phase would be random during fixation but undergo a reset at the time of the saccade (either onset or offset; onset pictured here). Here we seek to distinguish between these possibilities by analyzing the spectral content of ear canal recordings on individual trials.

Here, we assessed the pulse vs. phase resetting possibilities by evaluating microphone recordings obtained from ear canals of human participants with normal hearing while they performed a saccade task. We carried out spectral analyses and evaluated signal power and phase at the level of individual trials. We found evidence of both phase resetting and an increase in power in the 30-45 Hz frequency band at the time of saccades, with phase resetting present in 97% of participants but pulses present only in 40% of the subject pool. While some of the disparity between pulse vs phase-resetting in individual subjects could stem from possible differences in statistical sensitivity to these two different aspects of the waveform, not all of the disparity can be explained in this fashion: there were also differences in the relative timing of phase resetting vs power changes, and some subjects actually showed decreases in power at the time of the saccade.

The functional implications of the presence/prevalence of phase resetting are considerable. Broadly speaking, this observation releases certain constraints on our understanding of the mechanism(s) underlying EMREOs and adds new ones. The chief “released constraint” is that the underlying mechanism may not simply be creating a 30-40 Hz signal per se (a potential difficulty for various candidate mechanisms) but phase resetting at saccade onset/offset also contributes to the EMREO signal. Indeed, in some subjects, phase resetting alone appeared sufficient to educe EMREOs. The chief new direction is that we now need to understand not only how pulses of sound might be generated but also the mechanics of how such phase resetting might occur, as well as the respective contributions of actuators such as the middle ear muscles and outer hair cells. Ultimately, both pulsatile sound and phase resetting may be fingerprints of an underlying process, one that is theorized to facilitate adjusting how the auditory periphery responds to incoming sound in an eye-position dependent fashion. The results also provide a potential tie to previous studies showing oscillatory signals and/or phase resetting in the auditory pathway of the brain (Fu et al., 2004; Kayser, Petkov, & Logothetis, 2008; O’Connell et al., 2020; Obleser & Kayser, 2019; Schroeder, Lakatos, Kajikawa, Partan, & Puce, 2008) as well as studies that have identified rhythmicity (Dragicevic, Marcenaro, Navarrete, Robles, & Delano, 2019; Gehmacher et al., 2022; Kohler, Demarchi, & Weisz, 2021) and attention-related phase contrasts of ongoing otoacoustic activity in the cochlea (Köhler & Weisz, 2023).

## METHODS

Thirty human research participants (20 female, 10 male, age range 19-54 years, mean = 27.5 years) took part in the study. All procedures involving human subjects were approved by Duke University’s Campus Institutional Review Board. Subjects had clinically normal hearing (defined as pure tone thresholds less than 25 dB HL for 500, 1000, 2000 and 4000 Hz), tympanometry, and acoustic reflex thresholds (conventional 226 Hz probe tone). Methods for recording EMREOs are described in detail in previous publications (e.g., Lovich et al, PNAS, 2023; King et al, Hear Res, 2023). Microphone data (ER10B+) and eye tracking (EyeLink 1000) were recorded during performance of a saccade task with a central fixation target and saccade targets that were arrayed in a grid (Figure 2). Simultaneous recordings were made in both ears using ER10B+ microphones at 48 kHz digital sampling rate and then down sampled to 2000 Hz, providing a frequency range of 0-1000 Hz for analysis of oscillatory energy. As described in Lovich et al PNAS 2023, microphone data were synchronized to saccade onset or offset, with saccade onset defined as the first peak of the third derivative (jerk) and offset defined as the second peak. Data were smoothed prior to each differentiation using a 7 ms-long lowpass discrete filter to reduce the effects of noise and quantization error.

**Figure 2:**
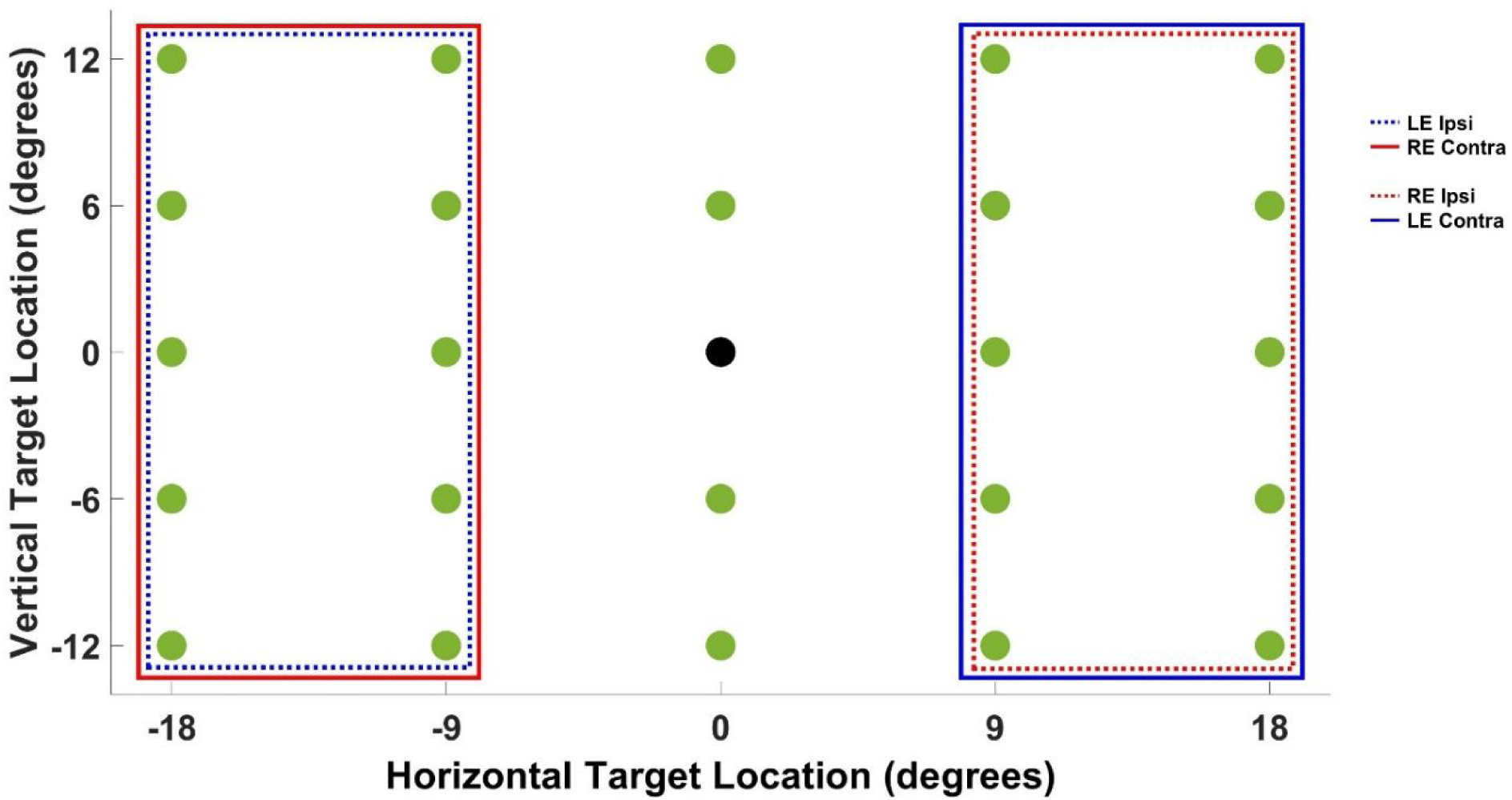
Schematic of grid task design and analysis. Each trial began with center fixation (black dot) followed by a saccade to one of twenty-four targets (green dots). We pooled the data into leftward saccades and rightward saccades (i.e. pooling across the vertical dimension, omitting the straight up and down targets) and by recording ear (left, right): LE-ipsi (left ear recordings during saccades to ipsilateral targets, dotted blue line), RE-contra (right ear recordings during saccades to contralateral targets, solid red line), RE-ipsi (right ear recordings during saccades to ipsilateral targets, dotted red line), LE-contra (left ear recordings during saccades to contralateral targets, solid blue line). Thus, the same group of targets were contralateral or ipsilateral, depending on recording ear.

Trials could be excluded based on either saccade performance or microphone noise. The exclusion criteria used for eye tracking were as follows: 1) presence of a sequence of two or more saccades to achieve the target; 2) eye tracking signal drop out during the trial (e.g., due to blinks); 3) slow or drifting non-saccadic eye movement 4) excessive saccade curvature of more than 4.5° (subtended angle); 5) inadequate duration (<200 ms) of steady fixation before and after the saccade. For microphone noise, trials were excluded if any microphone value on that trial exceeded 4 standard deviations above/below the mean microphone value across all trials for that session.

We verified that all subjects exhibited an EMREO using our previously described regression method (Gruters et al., 2018; C. D. King et al., 2023; Lovich et al., 2023): we fit the microphone data for each ear across time as follows:

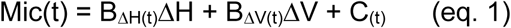

where ΔH and ΔV refer to the target locations in the horizontal and vertical dimensions, and the coefficients B_ΔH(t)_ B_ΔV(t)_ are time-varying coefficients designating the weighting of those factors. The constant term C_(t)_ is included to allow for any asymmetries in leftward vs rightward (or upward vs downward) in the signal. We evaluated the number of time points for which the regression fit was significant at p<0.025 in either the left or right ears during a time window −5 to 70 ms after saccade onset, matching our previous work (C. D. King et al., 2023). Chance values would be no more than 5% when combining across ears (significance would not be expected during zero crossings of this oscillatory signal, so in actuality the chance level is less than 5%). Observed values ranged from 19-99% across subjects (Supplementary Figure 1.

## RESULTS

EMREOs are most evident when averaged across at least a small number of trials, but in some subjects, the EMREO can occasionally be seen on individual trials. Figure 3 shows individual trial recordings from two participants. The set of trials in Figure 3A showed distinct “pulses” of sound on these trials, whereas those in Figure 3B showed phase resetting. The top panels show the individual saccade traces, involving an 18° rightward saccade. The lower panels show the associated ear canal recordings in the left ears, color coded to match the associated eye traces.

**Figure 3.**
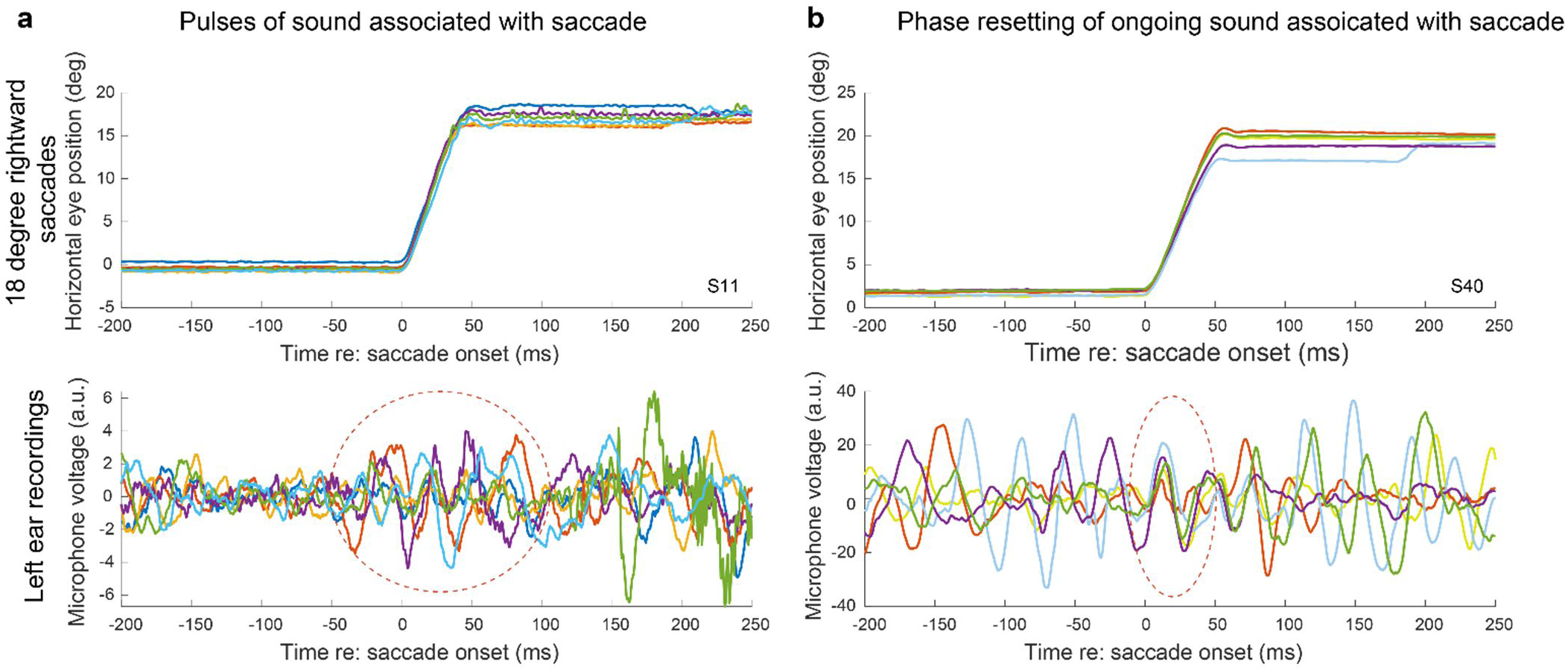
Example ear canal recordings on individual trials for two different participants (a, b) showing sound “pulses” (a) and phase resetting (b). Top panels show the eye traces for saccades to the 18 deg. leftward target (color indicates different trials). Bottom panels show the corresponding ear canal recordings during those trials (same color code). Dashed circle in (a) highlights a period of time aligned on saccade onset in which pulses of oscillation become apparent. Dashed circle in (b) highlights when ongoing oscillation comes into phase alignment across trials.

In Figure 3A, the dashed circle highlights a period of time in which the microphone signal begins oscillating at ∼25-30 Hz, starting on some trials as much as 50 ms before saccade onset. In contrast, from −150 to −50 ms before the saccade onset, the ear canal recording signal is relatively quiet. After the saccade, the signal remains elevated above that quiet pre-saccadic baseline period throughout the remainder of the trial, potentially related to the eye position component of the EMREO signal (Lovich et al., 2023) as the eyes are now fixating an eccentric position. Prior to −150 ms, additional oscillatory signals can be seen, potentially related to a prior saccade that brought the eyes to the fixation light at the beginning of the trial.

The pattern is different for the trials shown in Figure 3B, from a different participant but involving the same saccade target (18° right) and ear (left), i.e. contraversive saccades, again color coded such that the corresponding eye traces and ear canal recordings have matching colors. For this participant and set of trials, oscillatory energy is seen on at least some trials throughout the time window displayed, but these oscillations come into phase alignment at saccade onset (dashed circle) and appear to stay in alignment throughout the saccade.

To quantify these effects at the population level, however, several analysis choices needed to be made. We chose a time window of −25 to 75 with respect to either saccade onset or saccade offset, based on inspection of the trial-averaged EMREO waveforms of all individual subject ears (Supplementary Figure 2). Next, we measured the spectral power in this selected time window on these trial-averaged waveforms (Supplementary Figure 3). Microphone recordings during saccades to contralateral targets revealed a peak in spectral energy at ∼30 Hz while peak energy for ipsilateral targets was ∼40 Hz (5 Hz sampling resolution). To capture peak spectral energy for both contralateral and ipsilateral conditions, we chose a spectral band of 30-45 Hz for additional analysis. The peak spectral energy was similar for both onset and offset aligned analysis so we were able to use the same 30-45 Hz spectral band for both contexts of data analysis. Results were similar for left vs right ears, so subsequent analyses combine the data across all the ears in the sample (N=60).

With this spectral band chosen, we could then investigate the energy in this band on individual trials and across time as follows. First, we computed power and phase as a function of time in the EMREO frequency band (30-45 Hz) for individual trials (sliding 25 ms time bins with 50% overlap). Next, we combined the resulting trial-wise power and phase curves across ears for a given subject, yielding a subject-wise average for contraversive vs. ipsiversive saccades. Finally, we computed a grand average across the subjects to reveal the results at the population level. Figure 4 shows these grand average results for the data aligned to saccade onset and offset. The top row shows the power of the EMREO frequency band across time for the contraversive (blue lines) and ipsiversive (pink lines) saccades along with +/-1 SE, matching shaded areas. An increase in spectral power can be seen during the period of the EMREO signal (−25 to +75 ms) for both onset and offset aligned data. The bottom row plots the population average for phase. Changes in phase direction determined by saccade direction (contraversive vs ipsiversive) begin at the time of saccade onset/offset.

**Figure 4:**
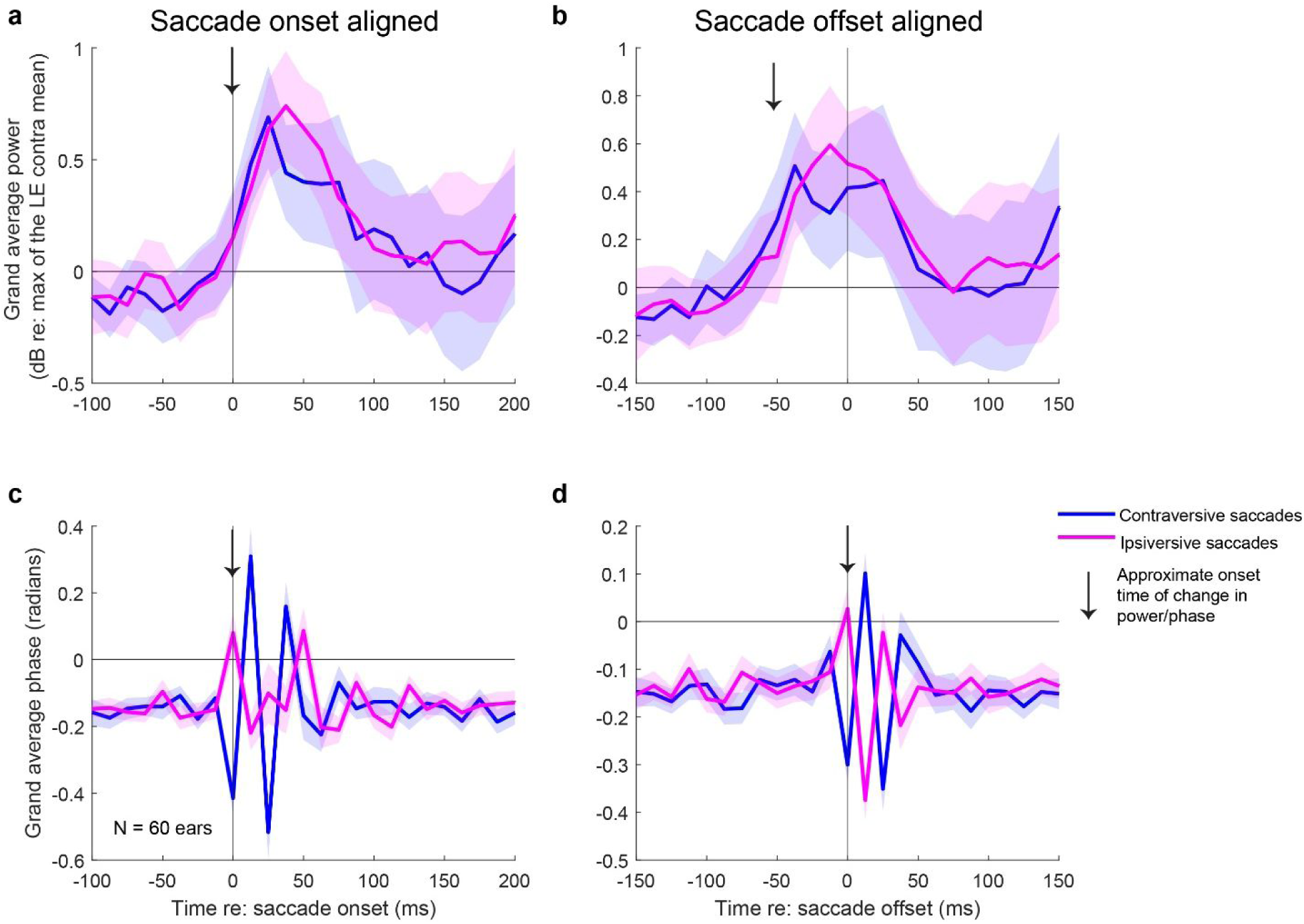
Power and phase of the EMREO frequency band aligned to saccade onset and offset. Power and phase were first computed at the individual trial level, then averaged within subject ears. Panels a, b show grand averages (N=30 subjects, 60 ears) of power in the 30-45 Hz frequency band, expressed in units of dB, relative to the maximum trial-averaged power observed for each subject’s left ear during contraversive (rightward) saccades. Blue and pink lines depict the signals measured during contraversive and ipsiversive saccades, respectively. Shaded areas are +/-1 SE. There is an increase in spectral power following saccade onset (a). When aligned to saccade offset (b), the increase in spectral power occurs prior to the end of the saccade (as expected since it is during the observed time window for the EMREO) and remains above steady fixation levels for approximately 50 ms after saccade offset. The bottom row are grand averages of phase. There is a clear change in phase at saccade onset (c) and offset (d), reflecting the phase reversal for ipsiversive vs contraversive saccades seen in time waveforms of the EMREO signal. The time courses of the power and phase changes are approximately the same as each other when aligned on saccade onset, but show a temporal disparity when aligned on saccade offset, as can be seen by the arrows which mark the approximate time of onset of the power increases (a, b) and phase changes (c, d). See supplementary Figures 4 and 5 for individual subject power and phase data.

The relative timing of the power vs phase effects is slightly different. For saccade onset, a phase signal is first evident in the bin at time 0 (arrow), with the power signal beginning around then (arrow) and peaking closer to the 25 ms time period. For saccade offset, the phase signal is again first evident in the bin at time 0 (arrow), whereas the power signal rises about 50 ms earlier (arrow) and is at a plateau at saccade offset. Thus, there is a timing discrepancy between these two measures that suggests a dissociation in the underlying cause.

In light of the individual differences evident in Figure 3, in which pulses vs. phase resetting appeared to be more or less prevalent in different participants, we next tested for statistical significance within individual participants. To test for significant pulses, we compared the spectral power in the EMREO frequency band during each saccade compared to a period of pre-saccade fixation using a one tailed pairwise ttest (paired saccade and pre-saccade periods for each trial). The time windows were selected based on visual inspection of the saccade onset-aligned grand average traces in Figure 4A, and were −100 to 0 ms before saccade onset for pre-saccadic fixation baseline and 12.5 to 62.5 ms after saccade onset for the saccade period. We conducted this test separately for contraversive and ipsiversive saccades, but we combined the data across ears. If the test was significant at the p<0.025 level for either contraversive or ipsiversive saccades, we considered that subject showed a significant “pulse” (the p<0.025 criterion provides for Bonferonni correction and thus, in net, this test can be assumed to have a false positive rate corresponding to p<0.05 or 5%). N= 12 of 30 participants (40%) showed a significant increase in spectral power in the EMREO frequency band after saccade onset by these criteria.

To test for the statistical significance of phase-resetting, we used a different method. Rather than conduct an analysis at the level of individual trials, we carried out the analysis across the time bins of the trial-averaged phase plots (i.e. the traces shown for each individual in Supplementary Figure 5), following a method commonly used to ascertain response latency in neural responses from peristimulus time histograms (Maier & Groh, 2010). To do this, we calculated the mean and standard deviation of the phase bins in a baseline period of –187.5 to - 25 ms re: saccade onset (phase values in 14 12.5 ms bins), again pooling across ears but keeping contraversive and ipsiversive saccades separate. We then took the maximum value of the phase observed during a time window (−12.5 to 50 ms re: saccade onset, five 12.5 ms time bins). Again, these time windows were chosen based on the grand averages shown in Figure 4C. If this maximum phase value exceeded 3.1standard deviations above the baseline period for either contraversive or ipsiversive saccade directions, the subject was considered as showing significant phase resetting. This criterion level was set based on Monte Carlo simulations to set the false positive rate for each saccade direction at 2.5% and thus the combined false positive rate across saccade directions at 5% (see legend to Table 1 for details).

**Table 1.** Relationship between statistical significance for “pulse” vs phase-resetting analyses. For the phase resetting analysis, a Monte Carlo simulation was conducted to set the criterion for significance as follows: simulations in which 14 data points were randomly drawn from a normal distribution, to simulate the baseline, and then 5 data points were randomly drawn from the same distribution to simulate the saccade windows. We ran this simulation 100,000 times to set the criterion such that only 2.5% of the simulated saccade-related maximum phases would exceed the simulated baseline by chance. The 2.5% criterion was applied for each saccade, so that across the two saccade directions, the chance rate would be 5%.

| sig. =<br>p<0.05 | Saccade onset aligned |  |  |  |  |  |
| --- | --- | --- | --- | --- | --- | --- |
|  | Phase sig. |  | Phase n.s. |  | Total |  |
|  | N | % | N | % | N | % |
| Pulse sig. | 12 | 40% | 0 | 0% | 12 | 40% |
| Pulse n.s. | 17 | 57% | 1 | 3% | 18 | 60% |
| **Total** | 29 | 97% | 1 | 3% | 30 | 100% |

Using this criterion, N= 29 of 30 participants showed phase resetting at saccade onset – all of the participants who showed significant increases in spectral power, plus 17 more who showed significance *only* for the phase (Table 1). Figure 5 shows the individual traces for each subject, with green indicating that the particular trace reached statistical significance, and gray if not. The differences between the phase-only group (blue box) vs the pulse-and-phase group (orange box) are clear: the phase signal is present in all cases, but a distinct pulse is evident only for the pulse-and-phase group. While there was one subject for whom phase did not reach significance (gray box), this may reflect the strictness of the criteria for significance rather than a true absence of phase resetting.

**Figure 5.**
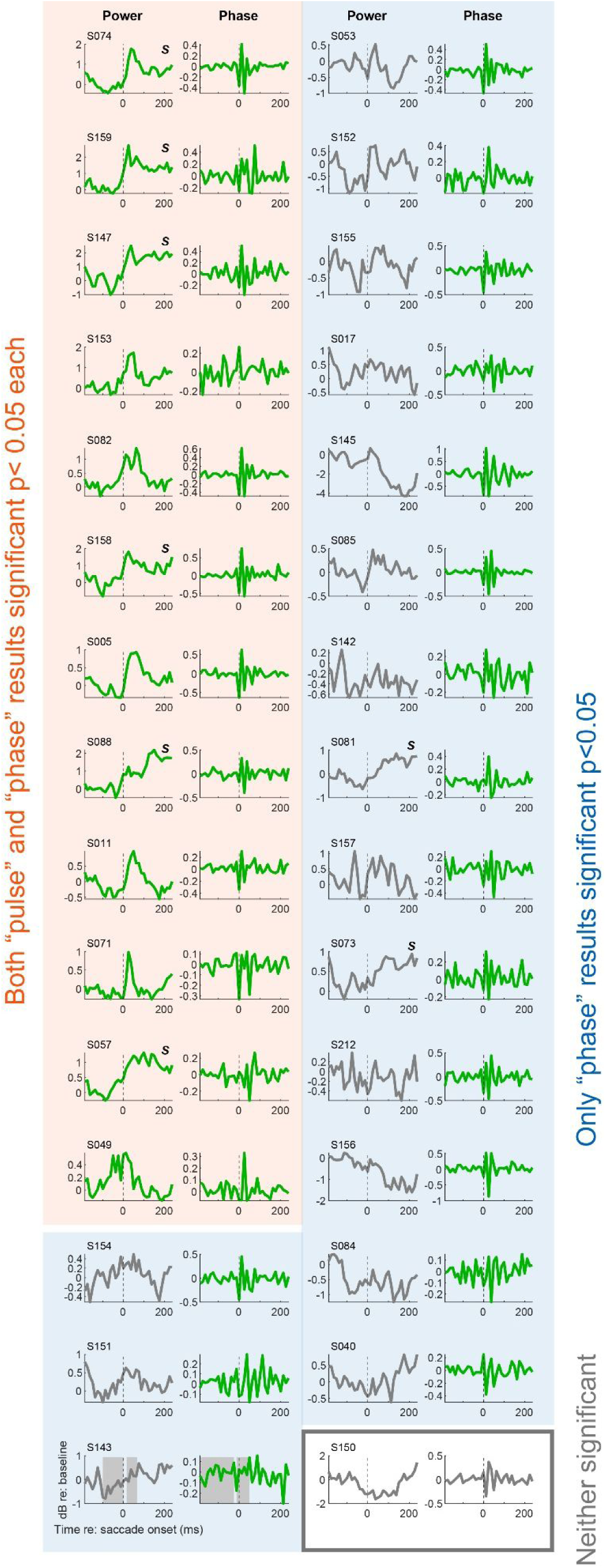
Spectral power and phase plots for each subject, grouped according to whether the change in power and/or phase reached statistical significance (green) or not (gray) for that subject. The orange box on the left shows the 15 subjects for whom both power and phase tests reached significance (p<0.025 for either contraversive or ipsiversive saccades, for a corrected p<0.05 within each type of analysis. The blue box groups the 13 subjects (43% of 30) for whom only the phase test reached significance. The gray box encompasses the subject for which neither test reached significance. Shaded areas in the lower left panels illustrate the time windows for analysis. The letter “S” in the top right for certain subjects indicates those mentioned in the main text as likely showing sustained or step-like increases in power during eccentric fixation. Contraversive and ipsiversive data were combined by averaging. The sign of the ipsilateral phase data was inverted before averaging.

Scrutiny of individual subjects reveals additional qualitative findings of potential interest. First, about 6 of the 13 subjects that showed statistically significant increases in power in the designated window comparing the saccade to baseline actually showed a step-like increase in power that persisted during the period of steady eccentric fixation after the saccade (S074, S159, S147, S158, S088, S057; see also 2 subjects who did not show significance for a power increase during the saccade also appeared to show this slightly later step-like power increase -- S081, S073).

Second, while the phase-resetting-only group may include some cases in which there is indeed a power increase at the time of the saccade, but which was too weak or too noisy to pass statistical criteria, there are also cases in which the power actually decreases around the time of the saccade (e.g. S156).

Finally, side-by-side comparison of the power vs phase curves reveals many examples of timing/duration differences, suggesting that the two metrics can be dissociable. For example, S074 shows a pulse + sustained step in power, but the phase curves show an oscillation that is more limited in duration.

## SUMMARY AND CONCLUSIONS

The main finding of this paper is that phase resetting appears to be widely evident when EMREOs are subjected to Fourier analysis at the trial level. Consistent with previous findings, phase changes were evident when time locked to both saccade onset and saccade offset. Indeed, the phase signal extends after the offset of the saccade for another 50+ ms. (Gruters et al., 2018; Cynthia D King et al., 2026). The phase change at saccade onset was observed in 97% of subjects (29 of 30). In contrast, saccade-related increases in sound energy were more variable across subjects than phase resetting, but were still present in 40% of subjects (12 of 30) at saccade onset. These power increases appeared to consist not only of short-duration pulses but also in some cases of “steps” that are sustained during eccentric fixation. In subjects that showed only a statistically significant phase resetting, the power signals showed variation: some subjects might have power increases that merely fell short of reaching statistical significance, but others actually showed a decrease in power around the time of the saccade.

As noted in the introduction, the pulse vs. phase alignment hypotheses are not mutually exclusive. Indeed, one might expect that if the underlying mechanism were to produce pulses at the time of the saccade, and these pulses were to all have a consistent phase, then both the power and the phase outputs of the Fourier analysis would show a significant association with saccades. (If there were pulses whose phase was random, then there would not be a significant phase signal, but neither would the trial-averaged waveform in the time domain have shown a significant signal – so the pulse-with-random-phase possibility could be eliminated based on previous findings.)

Accordingly, the most interesting aspect of the findings are dissociations between phase and power signals. At the population level, there is a temporal dissociation when the data are aligned on saccade offset: the power signal is already elevated at saccade offset whereas the phase resetting is not evident until that moment. This is curious because phase resetting is evident at saccade onset (and is synchronized with the increase in power) when the data are aligned that way, but it does suggest that the two metrics reflect dissociable processes.

At the individual level, dissociations were also observed: about 57% of the subjects showed significant phase resetting but not significant increases in sound power at saccade onset. As noted above, a caveat is that this observation rests on combining statistical analyses that (a) derive from the same underlying numbers, and (b) may not have equal power/susceptibility to false positives vs negatives. Nevertheless, the pattern is compelling enough to warrant consideration of its implications for the purpose of the EMREOs and the underlying mechanism(s) that generate them.

These results suggest that the perceptual purpose of EMREOs is not likely to be about (or not solely about) “adding” a 30-45 Hz sound to incoming sounds, but rather that the observed average pulses of sound signal is the result of some underlying mechanical shift that causes a phase change in ongoing sound energy. This underlying mechanical process might adjust the tension on the eardrum, altering the gain and timing of incoming sound transmission through the middle ear as postulated by (Cho et al., 2023). The phases were opposite across the two ears and within the same ear for opposite saccade directions, indicating that the underlying operation produces a binaural difference cue that depends on whether the saccade is directed to the left or the right.

Early speculation about EMREOs was that they likely had to be generated by the middle ear muscles (Gruters et al., 2018), because it would appear difficult for outer hair cells to generate a signal that was so large (averaging 56 dB SPL) (Lovich et al., 2023) and at such a low frequency. While it may or may not be possible for OHCs to expand/contract in concert at a low frequency to generate the EMREO wave, a role in phase adjustment might not require as much force at a specific frequency band as previously assumed.

The importance of phase in EMREO generation suggests that EMREOs may be important for more than one functional process. While we have focused chiefly on its potential role in adjusting how the auditory periphery responds to incoming sound in an eye-position dependent fashion, an additional possibility is that EMREOs contribute to temporal alignment of visual and auditory processing – time-stamping auditory processing to coordinate with the arrival of a fresh image on the retina. Indeed, several studies have suggested that eye movements modulate excitability in auditory cortex (Leszczynski et al., 2023; O’Connell et al., 2020), There is also evidence of oscillatory signals in the nervous system, including both central (Fu et al., 2004; Kayser et al., 2008; O’Connell et al., 2020; Obleser & Kayser, 2019; Schroeder et al., 2008) and peripheral auditory areas (cochlea/auditory nerve) (Dragicevic et al., 2019; Gehmacher et al., 2022; Kohler et al., 2021; Köhler & Weisz, 2023). Such oscillatory activity has been linked to attentional state, and phase resetting has been reported (Kayser, 2009; Lakatos et al., 2009). Most compelling is recent evidence that ongoing otoacoustic activity shows phase inversions across the two ears when attention is directed to one side vs. the other (Köhler & Weisz, 2023) Future work will be needed to tease apart the relative contributions of EMREOs to spatial and temporal coordination between the visual and auditory systems, as well as the relationship to these observations of oscillations in the central nervous system and their connection to attention.

Beyond the specific relationship to eye movements, this finding opens up new avenues for study in the mechanics of the ear and how active forces contribute. Phase resetting and realignment is commonly considered in the context of oscillations in the brain as noted above (for review, see (Canavier, 2015)), but to our knowledge has not yet been considered in ear mechanics.

## ACKNOWLEDGMENTS

This work was supported by NIH grant DC020363 to JMG and CDK. We thank Stephanie N. Lovich and Dr. Stephen Eliades for thoughtful comments on the analysis and/or manuscript.

## SUPPLEMENTARY FIGURES

**Supplementary Figure 1.**
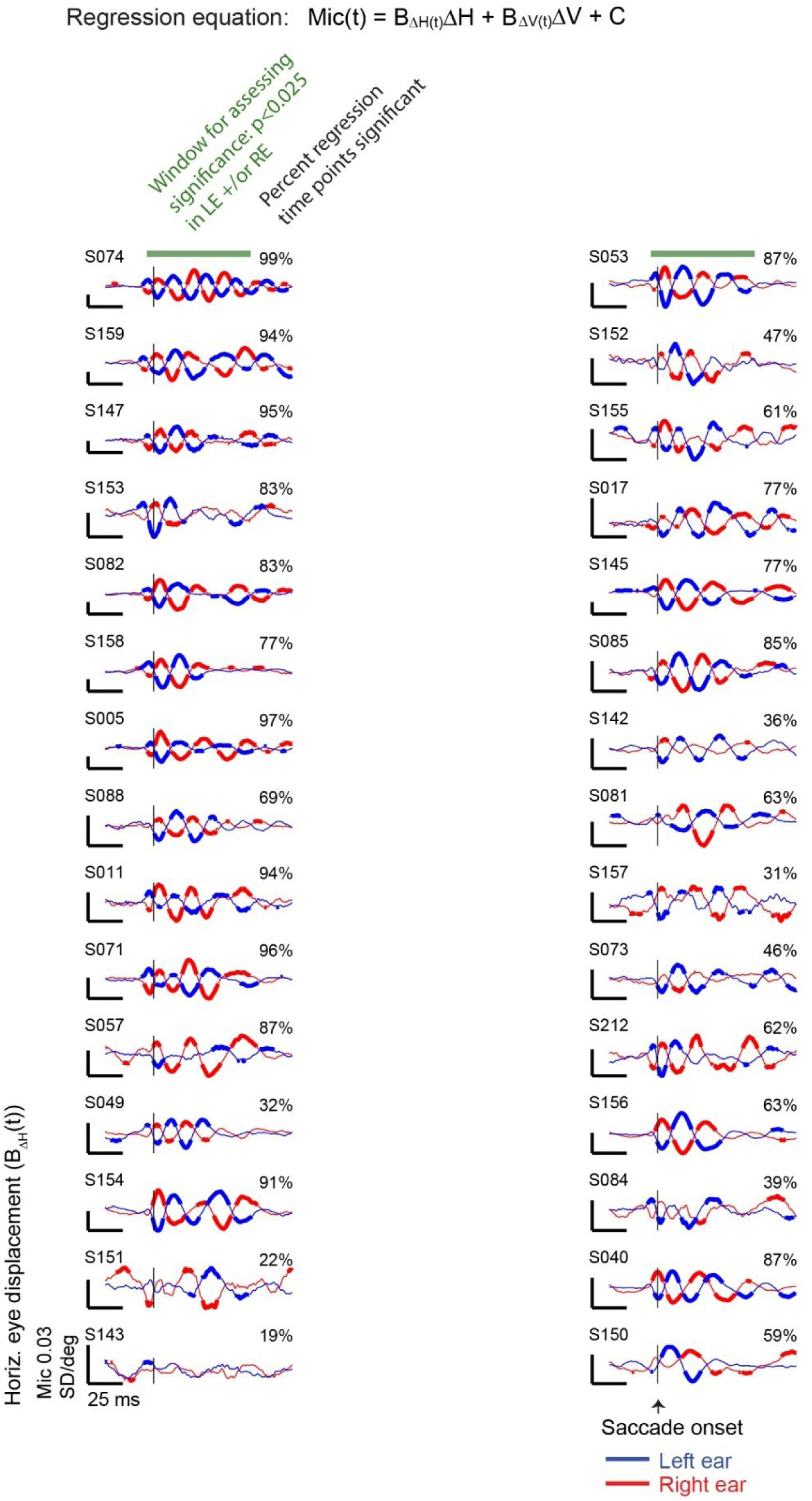
Assessment of statistical significance of the EMREO in individual subjects. Following our previous methods (C. D. King et al., 2023), we used regression to first establish that each of the included participants exhibited a measurable EMREO. The regression equation includes terms for both horizontal and vertical dimensions as illustrated. Each panel shows the horizontal coefficient for each subject for the left (blue) and right (red) ears. Each participant’s microphone data was Z-scored relative to their own baseline assessed −150 to −120 ms before the saccade onset, so the y-axis units are expressed as standard-deviation-per-degree. The percentage values are the percent of the time points that the overall regression fit was statistically significant during the time window −5 ms to 70 ms with respect to saccade onset (green bar) in either ear at p<0.025, for an aggregate false positive rate of 5%. Thick portions of the red and blue curves show periods of time when the 95% confidence intervals of the horizontal regression coefficient differed from zero.

**Supplementary Figure 2:**
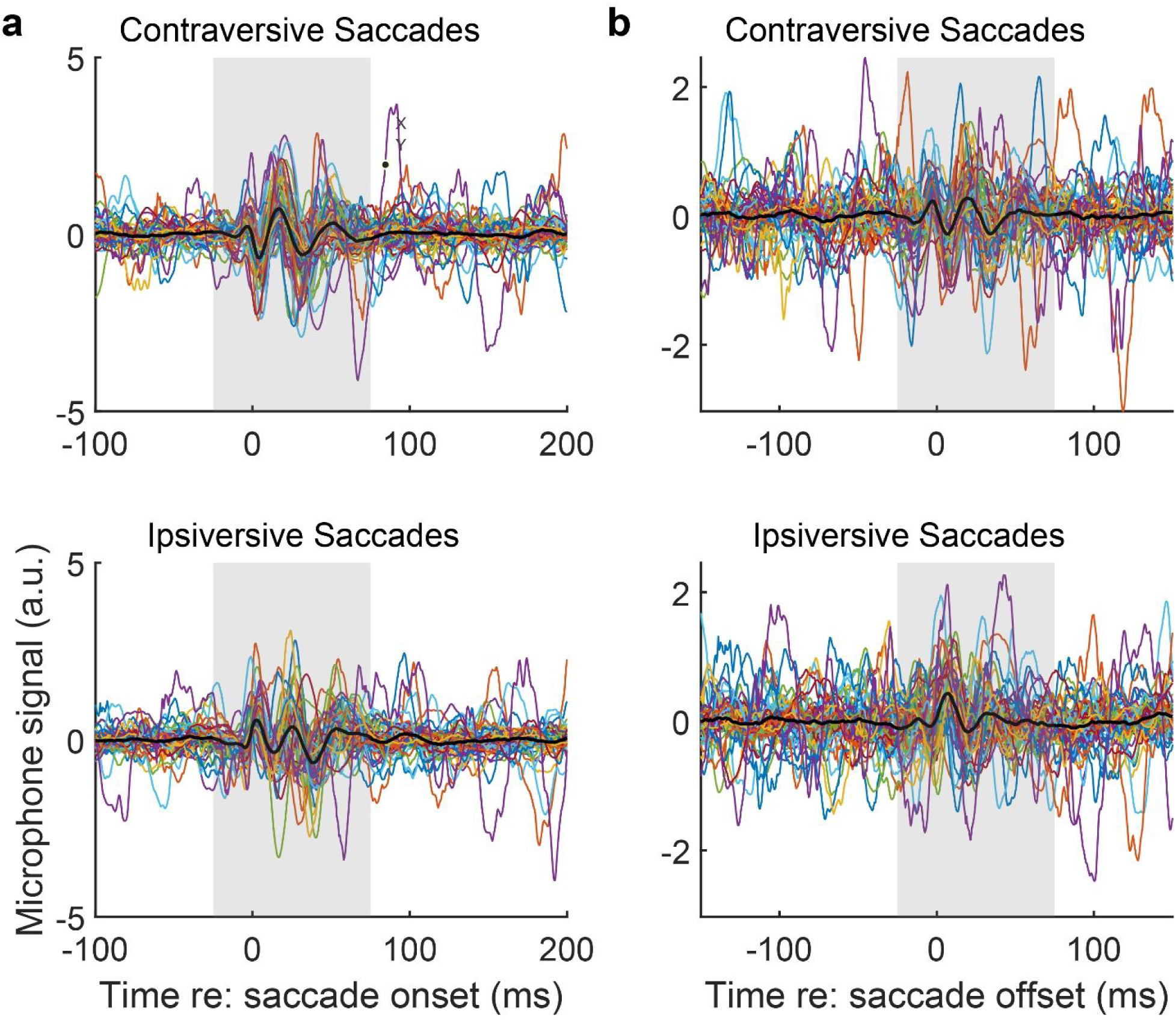
Individual ear average waveforms aligned to (a) saccade onset and (b) saccade offset. (a) Average waveforms aligned on saccade onset for all ears (colored lines, n=60), together with the grand average across ears (thick black line), grouped by contraversive and ipsiversive saccades. The purpose of this plot was to choose a time window to measure the overall spectral content around the time of the saccade. We chose a window of - 25 to + 75 ms (gray shaded) to fully capture the entire EMREO waveform during saccadic eye movement. As you can see, this window contains the full set of EMREO cycles in the grand average waveform (black line). (b) Same as (a) but aligned to saccade offset. The duration of the EMREO signal following saccade offset is similar to that seen for onset.

**Supplementary Figure 3:**
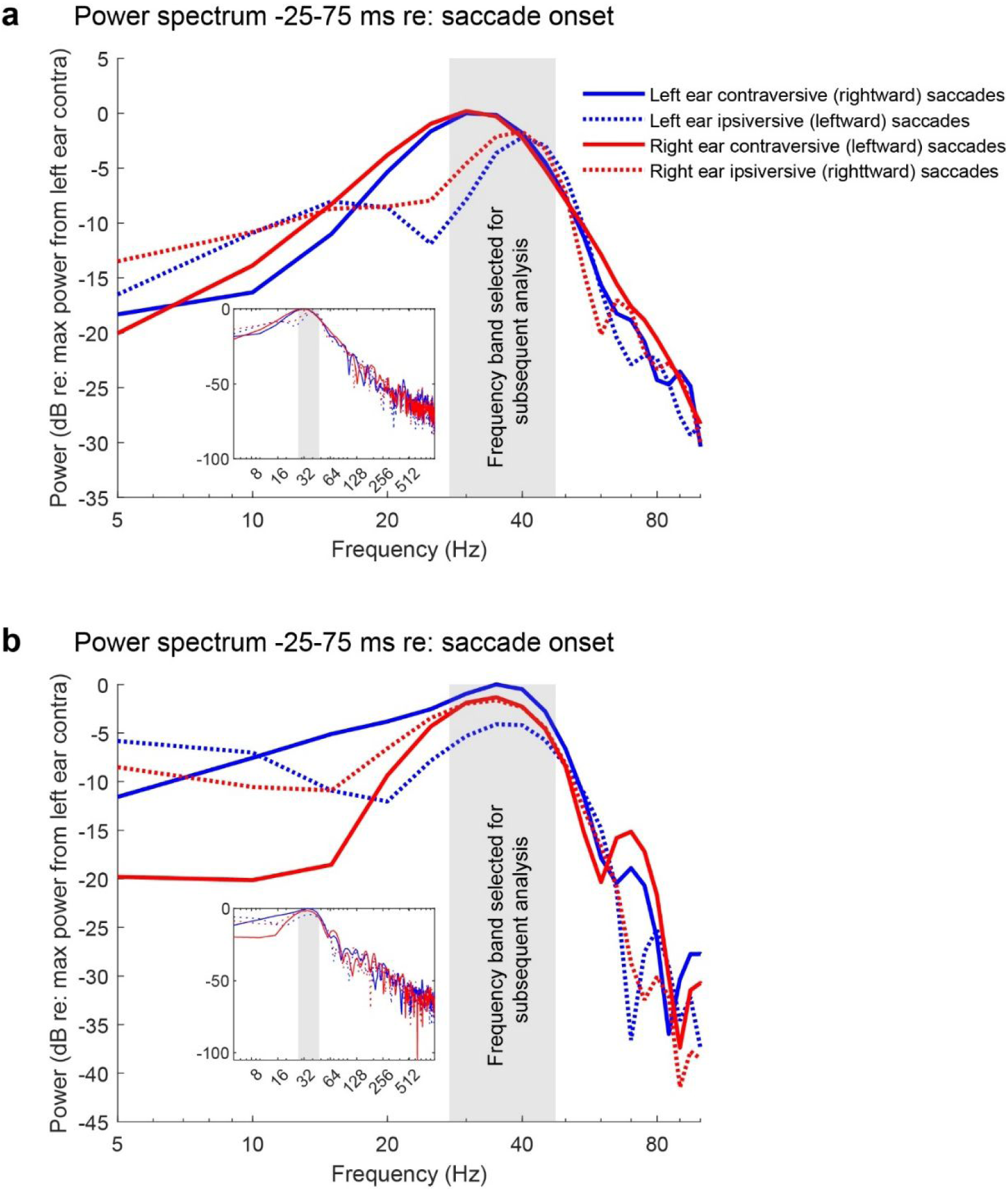
Average power spectrum for the EMREO time window for saccade onset (a) or offset (b). Shown here are the power spectrums of the grand average microphone recordings during the EMREO time window when aligned to saccade onset. The overall power spectra differed slightly depending on whether the saccades were contraversive (solid lines) vs ipsiversive (dashed lines) with respect to the recorded ear (right ear = red, left ear = blue), but were similar across ears when saccade direction was taken into account (red and blue curves are similar to each other). Accordingly, we selected a frequency band for analysis (30-45 Hz inclusive, gray box) that was wide enough to capture the peaks for both ipsiversive and contraversive saccades. We continued to separate data according to saccade direction but pooled across ears for subsequent analyses.

**Supplementary Figure 4.**
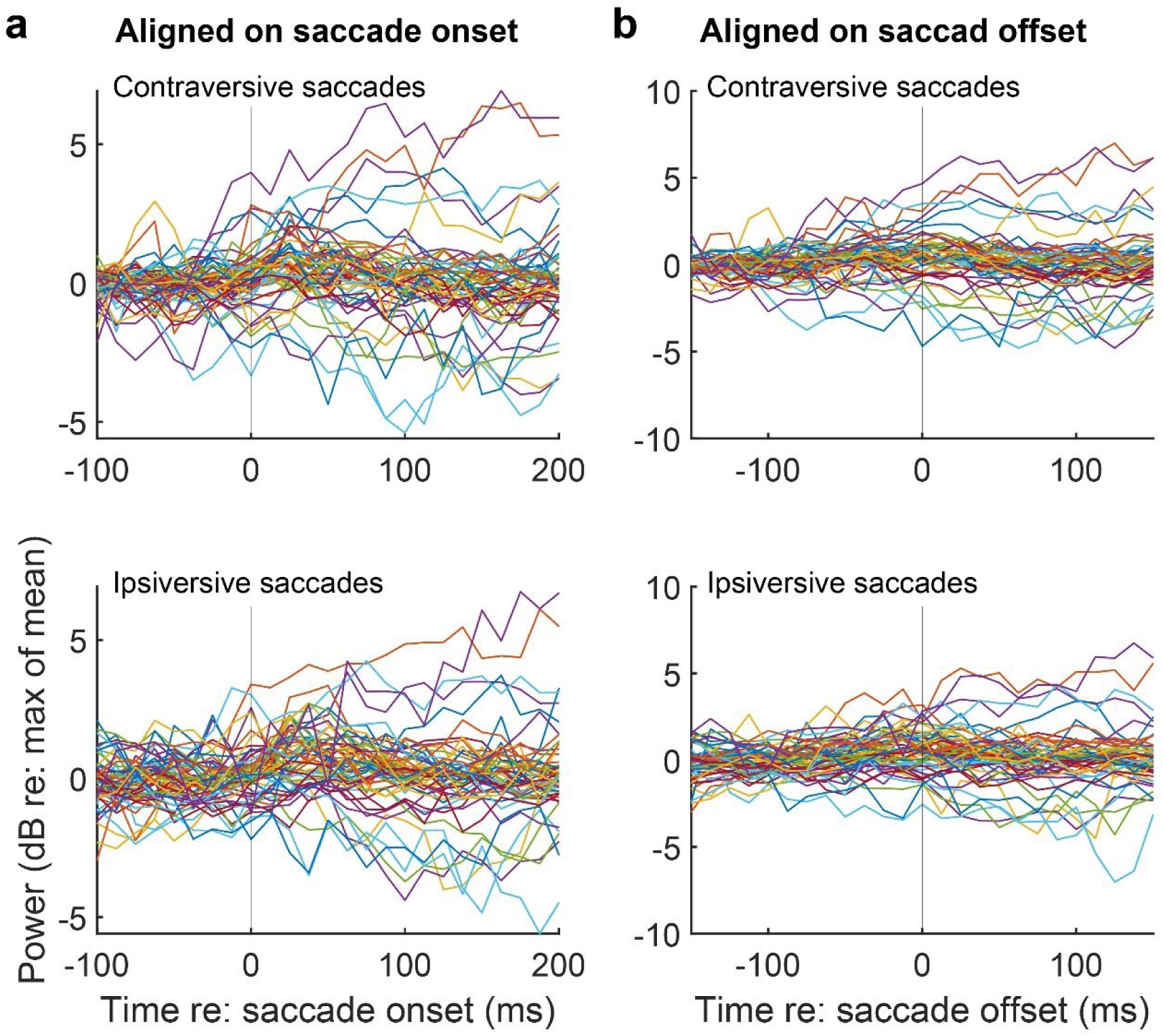
Spectral power for each ear (N=60). Plots of spectral power for the EMREO frequency band across time for contraversively and ipsiversively grouped data aligned to saccade onset (a) and offset (b). The increase in spectral power seen during the time of the saccade for the group average (*Figure 4*, first row) is not consistently present in all subjects.

**Supplementary Figure 5.**
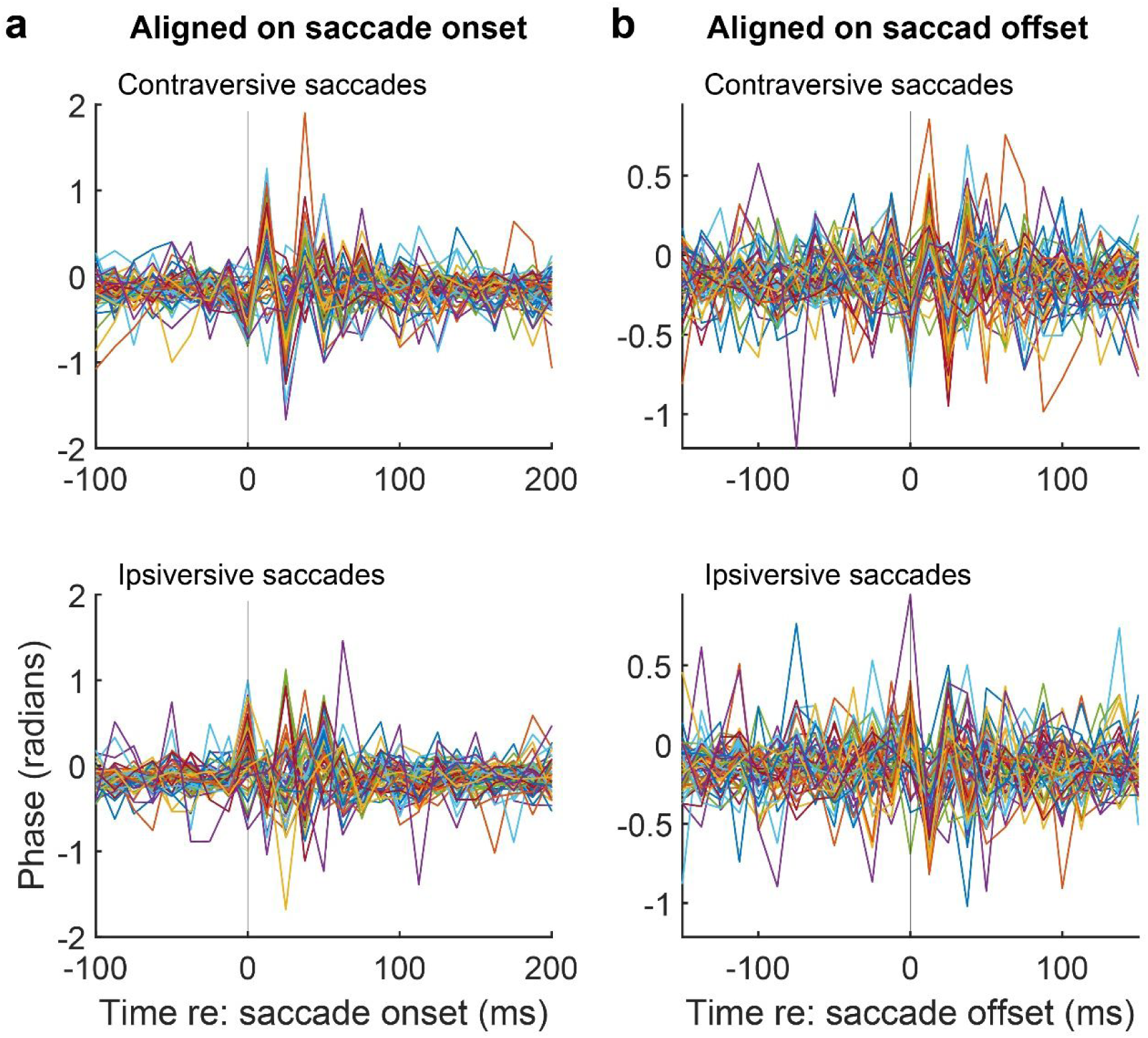
Signal phase in each ear. As before, data for the analysis groups based on saccade direction across all ears are presented when aligned to saccade onset (a) and offset (b). Each trace represents the average waveform of one ear (N=60). There is a very distinct, easily discernable signal that occurs at both saccade onset and saccade offset. While a distinct pattern for the population is response is evident, there are latency and amplitude variations across ears.

